# Phylogenetic evidence of headful packaging strategy in gene transfer agents

**DOI:** 10.1101/2020.10.08.331884

**Authors:** Emma Esterman, Yuri I. Wolf, Roman Kogay, Eugene V. Koonin, Olga Zhaxybayeva

## Abstract

Gene transfer agents (GTAs) are virus-like particles encoded and produced by many bacteria and archaea. Unlike viruses, GTAs package fragments of the host genome instead of the genes that encode the components of the GTA itself. As a result of this non-specific DNA packaging, GTAs can transfer genes within bacterial and archaeal communities. GTAs clearly evolved from viruses and are thought to have been maintained in prokaryotic genomes due to the advantages associated with their DNA transfer capacity. The most-studied GTA is produced by the alphaproteobacterium *Rhodobacter capsulatus* (RcGTA), which packages random portions of the host genome at a lower DNA density than usually observed in tailed bacterial viruses. How the DNA packaging properties of RcGTA evolved from those of the ancestral virus remains unknown. To address this question, we reconstructed the evolutionary history of the large subunit of the terminase (TerL), a highly conserved enzyme used by viruses and GTAs to package DNA. We found that RcGTA-like TerLs grouped within viruses that employ the headful packaging strategy. Because distinct mechanisms of viral DNA packaging correspond to differences in the TerL amino acid sequence, our finding suggests that RcGTA evolved from a headful packaging virus. Headful packaging is the least sequence-specific mode of DNA packaging, which would facilitate the switch from packaging of the viral genome to packaging random pieces of the host genome during GTA evolution.

## Introduction

Gene transfer agents (GTAs) are virus-like particles encoded and produced by certain bacteria and archaea (reviewed most recently by Lang et al. (2017) and Grull et al. (2018)). Unlike viruses, GTAs package fragments of the host genome instead of the genes that encode the GTA itself (Hynes et al., 2012). When GTA particles infect another cell, they can transfer the encapsidated genetic material to the recipient (Hynes et al., 2012; McDaniel et al., 2010; Solioz et al., 1975). The genomic loci that encode GTAs resemble prophages, indicating that GTAs evolved from viral ancestors. Although the function of GTAs is not firmly established, the prevailing hypothesis is that GTAs are not defective prophages, but instead are agents of horizontal gene transfer that are maintained in prokaryotic genomes due to the advantages associated with gene exchange, particularly, in stressful conditions (Kogay et al., 2020; Lang et al., 2017).

The best-studied GTA is produced by the alphaproteobacterium *Rhodobacter capsulatus*, and will be referred to as RcGTA. Production of the RcGTA particles is triggered by environmental factors (Westbye et al., 2017a), occurs in a small fraction of the population (Fogg et al., 2012; Hynes et al., 2012), is regulated by host proteins (Ding et al., 2019; Fogg, 2019; Westbye et al., 2017b) and involves expression of genes that are found in at least five loci in the *R. capsulatus* genome (Hynes et al., 2016; Lang et al., 2017). The largest of these loci is a 17-gene “head-tail cluster” that encodes proteins involved in head-tail morphogenesis and DNA packaging (Lang et al., 2017). There are homologs of the RcGTA head-tail cluster genes in other alphaproteobacteria (Shakya et al., 2017), including several *Rhodobacterales* for which GTA production has been observed (Fu et al., 2010; Nagao et al., 2015). Additionally, homologs of head-tail cluster genes are present in numerous viruses and proviruses (Shakya et al., 2017).

The small size of RcGTA and the low density of its packaged DNA precludes the particle from accommodating all of the genes required for its production (Bárdy et al., 2020). Instead, RcGTA packages seemingly random portions of the host genome (Hynes et al., 2012). In double-stranded DNA viruses of the realm *Duplodnaviria*, genome packaging is mediated by the terminase and portal proteins (Fokine and Rossmann, 2014; Rao and Feiss, 2015). Viral terminases typically consist of large and small subunits (Rao and Feiss, 2008). The small subunit (TerS) binds to the DNA to be packaged and then recruits the large subunit (Casjens, 2011; Rao and Feiss, 2015). The large subunit (TerL), which consists of ATPase and nuclease domains, cuts the concatemeric viral DNA, translocates the DNA into the viral capsid with concomitant ATPase hydrolysis and, finally, cuts the DNA again to terminate packaging (Casjens, 2011; Rao and Feiss, 2015). Viruses evolved different strategies for packaging DNA into their capsids, and these strategies involve different classes of TerL (Casjens and Gilcrease, 2009). Some viruses employ a “headful” packaging strategy in which TerL packages DNA into the capsid until it is full, whereas other viral TerLs terminate packaging at specific sequences (Casjens and Gilcrease, 2009; Rao and Feiss, 2008).

Viral TerLs with the same packaging mechanism tend to form clades in phylogenetic trees (Casjens et al., 2005). However, due to the diversity of TerL sequences, the relationships among the different functional classes of TerLs are not well-resolved (Casjens and Gilcrease, 2009; Casjens et al., 2005; Merrill et al., 2016). Given its lack of sequence specificity, RcGTA is presumed to package fragments of the host DNA via the headful strategy (Casjens et al., 2005; Hynes et al., 2012). The RcGTA TerL and its alphaproteobacterial homologs formed a distinct group in previous phylogenies, but they did not cluster with or within the viral TerLs that are involved in headful packaging (Casjens and Gilcrease, 2009; Casjens et al., 2005). Therefore, these phylogenies did not provide evidence of a headful packaging strategy in alphaproteobacterial GTAs, in part, due to limited sequence data available at the time.

In this study, we conducted a comprehensive evolutionary analysis of TerL sequences to better resolve the phylogenetic relationship of alphaproteobacterial RcGTA-like TerLs and viral TerLs with known packaging strategies. The results support a headful packaging strategy for RcGTA and, by inference, the rest of the putative alphaproteobacterial GTAs. We also identified two amino acid substitutions that are conserved in the TerLs of the putative alphaproteobacterial GTAs and might be important for the DNA packaging properties of GTAs.

## Methods

### Retrieval and Sequence-based Clustering of Large Terminase Homologs

Eighteen profiles covering one or both (ATPase and nuclease) domains of TerL were retrieved from the NCBI Conserved Domains Database (CDD) (Lu et al., 2020) (accessed on December 11, 2018). Two TerL profiles for distinct families of bacterial and archaeal viruses were added from Philosof et al. (2017) and Yutin et al. (2018). These 20 profiles (**Supplementary Table S1**) were used as queries for PSI-BLAST searches (E-value threshold of 0.01, effective database size of 2 × 10^7^ sequences) (Altschul et al., 1997) against the NCBI non-redundant protein database (accessed in December 2018). Only the subject sequences that were taxonomically assigned to archaea, bacteria and viruses were retained.

Partial TerL sequences were removed by ensuring the presence of both an ATPase (N-terminal) and a nuclease (C-terminal) domain using the following criteria: sequences either had to align to >= 75% of a “full” TerL profile that includes both TerL domains or align to different TerL profiles over their N- and C-terminal domains. A sequence was considered to align over a specific domain if it met one of two conditions: 1) if the sequence matched >=75% of an N- or C-terminal domain-specific profile and had at least 35% of the protein length outside of the matched domain to contain the unmatched domain or 2) if a sequence aligned to just the N- or C-terminal portion of a “full” TerL profile and had at least 35% of the protein length outside of the matched domain to contain the unmatched domain.

The resulting 254,382 sequences were clustered using MMseqs2 (Steinegger and Söding, 2017) with a similarity threshold of 0.75. From each of the obtained 11,298 clusters, a representative sequence of median length was selected for subsequent analyses.

### Alignment of Representative Homologs and Filtering Out Partial Sequences

The representative TerL sequences were iteratively aligned and clustered using the approach described by Wolf et al. (2018). Briefly, the sequences were clustered with a similarity threshold of 0.5 using UCLUST (Edgar, 2010). The clustered sequences were aligned using MUSCLE (Edgar, 2004) and alignment sites that contained more than 67% gaps were temporarily removed. Pairwise similarity scores between cluster alignments were calculated using HHSEARCH (Söding, 2005), converted to a distance matrix and used to build a UPGMA tree (Sokal and Michener, 1958). All of the branches of the UPGMA tree above a depth threshold of 2.3 were used to guide progressive alignment of the clusters using HHALIGN (Söding, 2005). The removed sites were reinserted back into their original sequences after the profile-profile alignment. These alignment and clustering steps were repeated for 20 iterations, when 11,230 of the sequences formed one alignment.

Of the remaining 68 sequences that failed to align, two were clearly TerLs but contained inteins, which were manually removed. Five other sequences were also likely TerLs because they were longer than 300 amino acids, exhibited significant similarity to a TerL profile via CDD searches (E-value < 0.001), and contained recognizable Walker A motif and nuclease catalytic residues. The seven sequences were profile-aligned to the alignment of 11,230 sequences using more relaxed criteria (similarity threshold of 0.01 and UPGMA depth threshold of 6). The remaining 61 sequences did not meet these criteria and were discarded.

The new alignment of 11,237 sequences contained partial sequences that lacked a Walker A motif. To remove these, each sequence’s similarity was scored to the alignment’s consensus sequence using a BLOSUM62 substitution matrix and the score was compared to the score of sequences with 100% identity to the consensus sequence. Sequences with a score less than 10% of the perfect match score were removed. Then, the alignment was used as a PSSM in a PSI-BLAST search against all of the sequences within the alignment. Only the sequences that matched to >= 75% of the PSSM were retained. The sections of the 11,060 sequences that passed this criterion were extracted and re-aligned using the above-described iterative alignment procedure with a clustering similarity threshold of 0.5 and UPGMA depth threshold of 2.3. After 23 iterations of alignment and clustering, 11,057 of the sequences aligned. The three sequences that did not align were longer than 300 amino acids, exhibited significant similarity to a TerL profile via CDD searches (E-value < 0.001), and contained a recognizable Walker A motif and nuclease catalytic residues. Therefore, they were retained and aligned to the main alignment using more relaxed parameters of a clustering similarity threshold of 0.01 and UPGMA depth threshold of 6.

### Alignment Trimming

The alignment of 11,060 sequences was trimmed to remove all columns with more than 50% gaps and less than 10% amino acid homogeneity. The homogeneity value of an alignment column was defined and calculated using the following procedure. For each of the ! = 11,060 sequences, column-based sequence weights 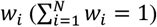 were assigned according to Henikoff and Henikoff (1994). The score of an alignment column against an amino acid *x* was calculated as 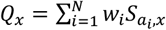, where *a_i_* is an amino acid in the *i*-th sequence and *S_a_i_,x_* is the BLOSUM62 substitution matrix score for a pair of amino acids *a_i_* and *x* (Henikoff and Henikoff, 1993). As the consensus amino acid of the column *c*, the amino acid with the highest score *Q_c_*, i.e. 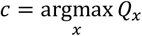, was selected. An expectation of the score of the given alignment column against a randomly selected amino acid *R* was calculated as *Q_R_* = ∑*_b_ f_b_Q_b_*, where *f_b_* is the vector of relative frequencies of amino acids (∑*_b_ f_b_* = 1, *b* ∈ {Ala.. Tyr}). The homogeneity of an alignment column was defined as 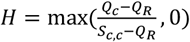. The homogeneity ranges from 0 (the alignment column score does not exceed the random expectation score *Q_R_*) to 1 (the alignment column score is equal to the maximum possible score *S_c,c_*).

### Reconstruction of Large Terminase Phylogeny

The trimmed alignment of 11,060 sequences was used to reconstruct an initial phylogenetic tree in FastTree v. 2.1.4 (Price et al., 2010) using the Whelan and Goldman (WAG) substitution model (Whelan and Goldman, 2001) and 20 gamma-distributed rate categories (Yang, 1994). The initial tree was used as a guide tree to refine our alignment, which in turn was used to reconstruct an improved tree. To this end, the sequences were divided into two sets, those that formed distinct groups on the tree and the remaining ones. The sequences that formed distinct groups were re-aligned using a clustering similarity threshold of 0.01 and UPGMA depth threshold of 2.3, whereas the other sequences were aligned using more stringent parameters (similarity threshold of 0.66 and UPGMA depth threshold of 1.3). These alignments were profile-aligned using a similarity threshold of 0.5 and UPGMA depth threshold of 2.3. Nine sequences did not join the main alignment and were discarded due to low scores against the consensus, calculated with a BLOSUM62 matrix as described above. The resulting alignment of 11,051 sequences was trimmed to remove all columns with more than 50% gaps and less than 10% amino acid homogeneity, as defined above. The final phylogenetic tree of 11,051 TerL homologs (**Figure 1**) was reconstructed using FastTree v. 2.1.4 (Price et al., 2010) with the WAG substitution model (Whelan and Goldman, 2001) and 20 gamma-distributed rate categories (Yang, 1994).

**Figure 1.**
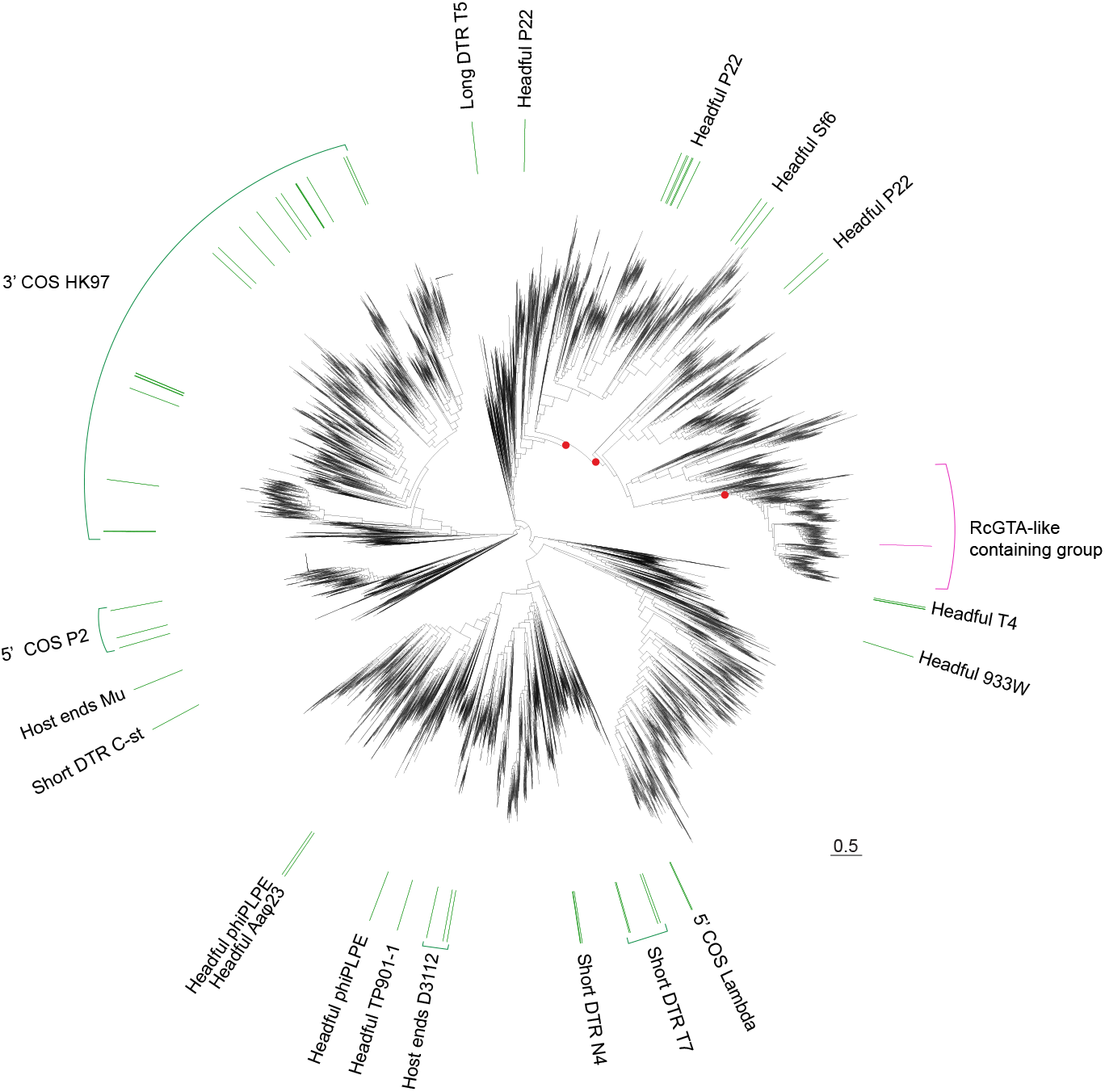
Phylogeny of TerLs from 11,051 viruses, prophages and GTAs. The TerL protein from RcGTA is denoted by a pink bar. The pink bracket outlines a subtree that contains RcGTA-like TerLs and is shown in detail in **Figure 2**. The green bars denote the viruses that have experimentally determined packaging strategies (**Supplementary Table S2**). The viruses with experimentally determined packaging strategies are labeled by the packaging strategy (Headful, Cohesive ends [COS], Direct Terminal Repeats [DTR], and Host Ends) followed by a prototype phage from that group (e.g. P22). Support values (aLRT) of > 75% denoted by red dots are only shown for a selection of branches relevant to grouping RcGTA-like TerLs within phages that employ a headful DNA packaging strategy. Scale bar, amino acid substitutions per site. Tree topology with all support values is available in NEWICK format in **Supplementary Dataset**. The patterns of this phylogeny are consistent with those in the phylogenies reconstructed using a more accurate maximum likelihood inference carried out using the IQ-TREE program (see **Supplementary Dataset)**; in particular, the RcGTA-like TerLs continue to form a well-supported group (100% of bootstrap samples), which forms a sister group to TerLs of phages that utilize a headful packaging strategy (75% of bootstrap samples).

### Identification of RcGTA-like Large Terminases

In the phylogenetic tree of TerLs (**Figure 1**), a group of 616 TerLs was labeled as the “RcGTA-like containing group.” This group includes 507 TerLs of the 526 RcGTA-like TerLs that were identified and curated by Kogay et al. (2019). Nineteen RcGTA-like TerLs from the dataset of Kogay et al. (2019) are absent in our dataset because they were not in GenBank at the time of our data collection (December 2018). The 616 TerLs were classified as either “RcGTA-like” or “virus-like” using a machine learning approach implemented in the GTA-Hunter program (Kogay et al., 2019).

### Examination of the Genomic Neighborhoods of Large Terminases

For the 604 TerLs within the “RcGTA-like containing” group (**Figure 1**) that originated from bacterial and archaeal genomes, the presence of 11 other RcGTA-like genes near the *terL* gene was examined. To this end, the RcGTA-like genes from the training set of Kogay et al. (2019) were used as queries in a BlastP search against the assemblies of the bacterial and archaeal genomes (E-value < 0.001; query and subject overlap by at least 60% of their length) (Altschul et al., 1997). The detected RcGTA gene homologs were classified as “RcGTA-like” or “virus-like” using GTA-Hunter (Kogay et al. 2019). The detected RcGTA gene homologs were also clustered into regions using DBSCAN, with a maximum distance cutoff of 8,000 bp between adjacent genes (Ester et al., 1996; Kogay et al., 2019). If a *terL* gene was embedded in a region containing at least 6 of the 11 RcGTA-like genes, it was classified as being in a “large RcGTA-like element.” If a *terL* gene was located in a region containing 1 to 5 RcGTA-like genes, it was classified as being in a “small RcGTA-like element.” The classification of the sequence represented in the phylogeny was assumed to be the same for the rest of the cluster members although this was not directly verified.

### Assignment of Packaging Strategy to Viral Large Terminases

A list of viruses with experimentally determined packaging mechanisms was compiled from the phylogenetic tree of Casjens and Gilcrease (2009). Viruses from the phylogenetic tree of Merrill et al. (2016) were also added, for most of which experimental evidence of the packaging mechanism is available. The dataset of the 252,614 TerL homologs was searched for these 87 viruses using TerL accession numbers provided by Merrill et al. (2016) and NCBI taxonomy IDs for the viruses from Casjens and Gilcrease (2009). Of the 87 viruses, 73 were present in our dataset (**Supplementary Table S2**). Due to the close sequence similarity, some of the 73 TerLs belong to the same MMSEQ clusters, and therefore are represented by 58 TerLs on our phylogeny of 11,051 TerLs (**Figure 1**). The 58 representative viruses for which the packaging mechanism was not known were assigned the mechanism of a virus from the same cluster with a known packaging strategy, under the assumption that the similarity of their TerL amino acid sequences is sufficient to imply the same packaging mechanism.

### Validation of the Reconstructed Phylogenetic Patterns with More Accurate Maximum Likelihood Analyses

To confirm that the phylogenetic relationships obtained from the FastTree program (Price et al., 2010) were not impacted by its limited tree search and optimization capabilities, additional phylogenetic trees were reconstructed using IQ-TREE v 1.6.7 (Nguyen et al., 2015) from two datasets subsampled from the 11,051 TerLs. The first dataset of 342 TerLs was constructed to broadly represent the TerL diversity (**Figure 1**). The dataset contains the 58 representative viral TerLs (described in **Assignment of Packaging Strategy to Viral Large Terminases** section), 50 TerLs randomly sampled from group 1 of the “RcGTA-like containing group” (**Figure 2**), all TerLs from group 2, 70 TerLs from group 3, 50 TerLs randomly sampled from the rest of the “RcGTA-like containing group,” and 100 randomly sampled TerLs from the rest of the whole TerL tree (**Figure 1**). The second dataset of 346 TerLs was constructed to represent well the TerLs from the “RcGTA-like containing group” (**Figure 1** and **Figure 2**). The dataset contains 12 representative viral TerLs with either P22 or Sf6-like headful packaging strategies, 70 TerLs randomly sampled from group 1, all TerLs from group 2, and 250 TerLs randomly sampled from group 3. For both datasets, the aligned sequences were retrieved from the trimmed alignment of 11,051 TerLs (see **Reconstruction of Large Terminase Phylogeny** section) and gap-only sites were removed. The optimal evolutionary model was selected using ModelFinder (Kalyaanamoorthy et al., 2017), as implemented in IQ-TREE. Support values for branches were calculated using ultrafast bootstrap approximation with 1000 replicates (Hoang et al., 2018), as implemented in IQ-TREE. The tree reconstructed from the second dataset was rooted with the headful P22 viral TerL that, out of the headful P22 viral TerLs in **Figure 1**, is the most distantly related to the “RcGTA-like containing group.”

**Figure 2.**
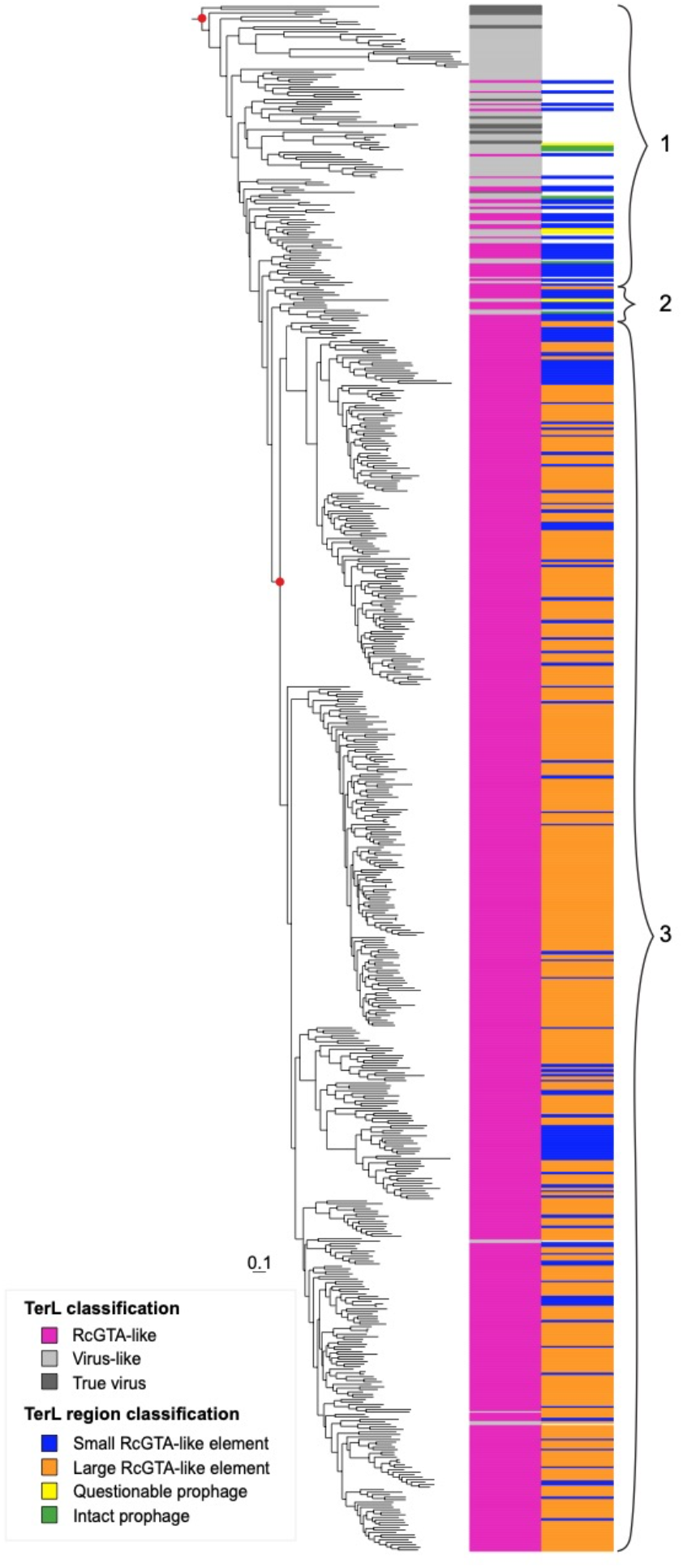
A subtree of the phylogeny shown on Figure 1 that contains RcGTA-like TerLs. Bars in the first column next to the branches of the subtree indicate whether the TerLs are from true viruses or, if they are found in bacterial or archaeal genomes, whether they were classified by GTA-Hunter as “RcGTA-like” or “Virus-like.” Bars in the second column denote whether the genomic neighborhood of the *terL* gene contains at least 6 RcGTA-like genes (“large RcGTA-like element”), between 1 and 5 RcGTA-like genes (“small RcGTA-like element”), a questionable prophage or an intact prophage. TerLs without a colored bar in the second column were not predicted as being in a prophage or RcGTA-like element. Group 1 includes mostly virus-like TerLs as well as TerLs from predicted prophages and true viruses. Group 2 contains a mixture of TerLs from a large element, small RcGTA-like elements and predicted prophages. Group 3 mostly contains RcGTA-like TerLs from “true” GTAs (large RcGTA-like elements). Red dots indicate two nodes that are relevant for the grouping of the entire subtree and for the TerLs in group 3, and have aLRT support values of > 75%. The patterns of this phylogeny are consistent with those in the phylogeny reconstructed using IQ-TREE (see **Supplementary Dataset)**; in particular, the node corresponding to group 3 has 96% bootstrap support. Scale bar, amino acid substitutions per site.

### Annotation of Prophages in Regions encoding Virus-like TerLs

To examine whether the bacterial TerLs classified as virus-like reside in prophages, prophages were predicted in the corresponding nucleotide genome sequences using PHASTER (Arndt et al., 2016, accessed in May 2020). The regions labeled as intact (score > 90) or questionable (score 70-90) prophages were retained, while the regions labeled as incomplete prophages (score < 70) were discarded. One region surrounding the virus-like TerL (accession WP_020474221) was predicted by PHASTER to be an intact prophage and was also classified as a small RcGTA-like element by GTA-Hunter, due to the presence of one RcGTA-like gene. The PHASTER prediction of this region as a prophage was considered to supersede the small RcGTA-like element classification.

### Detection of Conserved Sites that Differentiate RcGTA-Like and Virus-Like TerLs

The amino acid sequences of the 616 TerLs in the subtree shown on **Figure 2** were re-aligned using MAFFT-linsi v. 7.305 (Katoh and Standley, 2013). The alignment was scanned for sites conserved in more than 80% of the TerL homologs from group 1 (virus-like) or group 3 (RcGTA-like) (**Figure 2**). Of these detected sites, only the sites with at least 70% between-group difference in the relative abundance of the most conserved amino acid were retained.

The positions of the two identified sites relative to the known TerL structural domains were determined by searching CDD (Lu et al., 2020) with RcGTA TerL RefSeq record WP_031321187 as a query (database accessed on July 26, 2020). The conservation of the two detected sites within the “RcGTA-like containing group” was visualized using a subset of the 616 TerLs (**Figure 2**): 15 randomly sampled TerLs from group 1, all TerLs from group 2, 14 randomly sampled TerLs from group 3, and a representative TerL from the cluster that contains the RcGTA TerL. The aligned TerLs were retrieved from the alignment of 616 TerLs, and the gap-only alignment positions were removed. Secondary structure information was obtained via HHPred using RcGTA TerL as a query (Zimmermann et al., 2018).

The locations of the two sites were also visualized on a 3D structure of TerL from the *Shigella* phage Sf6 (PDB ID 4IDH) (Zhao et al., 2013) which, based on our phylogenetic inference, is the TerL most closely related to RcGTA-like TerLs for which a structure is available. The homologous positions of the substitutions in the *Shigella* phage Sf6 TerL were identified by aligning it to the RcGTA TerL using HHPred (Zimmermann et al., 2018). The visualization was carried out in PyMOL v 2.4 (Schrödinger, LLC, 2020).

## Results

### RcGTA-like TerLs belong within the group of TerLs of headful packaging phages

Of the 11,051 representative TerL homologs from bacteria, archaea and viruses, 616 are closely related to the RcGTA TerL and form a well-supported group in the phylogenetic tree (**Figure 1**). Of these 616 TerLs, twelve are encoded in viral genomes, whereas the remaining 604 are found in 601 bacterial and 3 archaeal genomes. Using a machine learning approach that relies on amino acid composition, we classified the 604 bacterial and archaeal TerL homologs as either “RcGTA-like” (527) or “virus-like” (77) (**Supplementary Table S3**). By mapping 73 TerLs with experimentally determined packaging strategies onto the phylogeny, we found that RcGTA-like TerLs fall, with strong support, within a group of headful packaging phages (**Figure 1**). Therefore, our phylogeny implies that the RcGTA-like TerLs evolved from a viral TerL that employed a headful packaging strategy and thus supports the hypothesis that RcGTA-like TerLs use a headful mechanism to package host DNA (Casjens et al., 2005; Hynes et al., 2012). Of the TerLs from the viruses with experimentally determined packaging strategies, *Enterobacteria* phage P22-like TerLs are the closest relatives of the RcGTA-like TerLs. This affinity contrasts the previous results, from analyses of much smaller data sets, according to which T4-like (Hynes et al., 2012; Lang and Beatty, 2000) or T7-like (Casjens et al., 2005) TerLs have been found to be most closely related to the RcGTA-like TerLs.

### Phylogenetic evidence of a single origin of GTA TerLs

Whereas the TerLs of viruses that employ headful DNA packaging specifically package the viral genome into the capsid, RcGTA TerL lacks sequence specificity and packages random segments of the bacterial genome (Hynes et al., 2012). To evaluate if non-specific DNA packaging evolved once or multiple times, we first sought to determine more accurately which of the RcGTA-like TerLs likely belong to *bona fide* GTAs. Shakya et al. (2017) hypothesized that genomic regions with a smaller number of recognizable RcGTA gene homologs are more likely to be prophages than GTAs because these regions tend to have a more virus-like GC content relative to their host, evolve faster and are more often associated with viral genes. Therefore, some of the TerLs classified as “RcGTA-like” might not belong to RcGTA-like elements in cases when the alphaproteobacterial genomes that contain these genes lack homologs of other RcGTA genes. Among the 527 RcGTA-like TerLs, we classified 391 as “large” (containing at least 6 RcGTA-like genes near the *terL* gene) and 136 as “small” (1-5 RcGTA-like genes) elements (**Figure 2, Supplementary Table S3**).

Within the subtree that contains RcGTA-like TerLs (**Figure 2**), all but one of the TerLs found in large elements form a well-supported clade (group 3 in **Figure 2**). The one TerL from a large element that falls outside this clade is a representative of a cluster of three TerLs that are found in the genomes of alphaproteobacteria *Zavarzinia compransoris* DSM 1231, *Zavarzinia* sp. HR-AS and *Oleomonas* sp. K1W22B-8. This TerL belongs to a narrow “transition zone” (group 2 in **Figure 2**) between the group 3 TerLs and the deepest branches of the subtree that include exclusively viral and “virus-like” sequences (group 1 in **Figure 2**). The transition zone also contains a mix of RcGTA-like TerLs from small elements and virus-like TerLs, including the TerL from the intact prophage predicted in the genome of a planctomycete *Zavarzinella formosa* DSM 19928. None of the TerLs within this transition zone come from functionally characterized viruses or GTAs. Thus, the phylogeny indicates that RcGTA-like TerLs likely evolved only once from a viral TerL, in an ancestor of group 3, by acquiring the capability to package DNA non-specifically. The positions of the TerLs from *Zavarzinia’s* and *Oleomonas’* putative GTAs and the *Zavarzinella* prophage could be explained by horizontal gene transfer, as previously documented in some instances for other RcGTA-like genes (Yang et al., 2017) and discussed in detail below.

### Viruses might mediate horizontal gene exchange of RcGTA-like genes

In addition to the above-discussed predicted prophage from *Zavarzinella formosa* DSM 19928, we identified two intact prophages in the genomes of the firmicute *Thermoactinomyces* sp. DSM 45892 and the alphaproteobacterium *Methylobacterium terrae* 17Sr1-28, and 16 viruses that encode TerLs that are phylogenetically most closely related to the RcGTA-like TerLs (**Tables 1 and 2**). Notably, the TerLs of these three prophages are even more closely related phylogenetically to the TerLs from large elements than the 16 viruses are (**Figure 2**), but whether they produce functional virions is unknown.

**Table 1.**
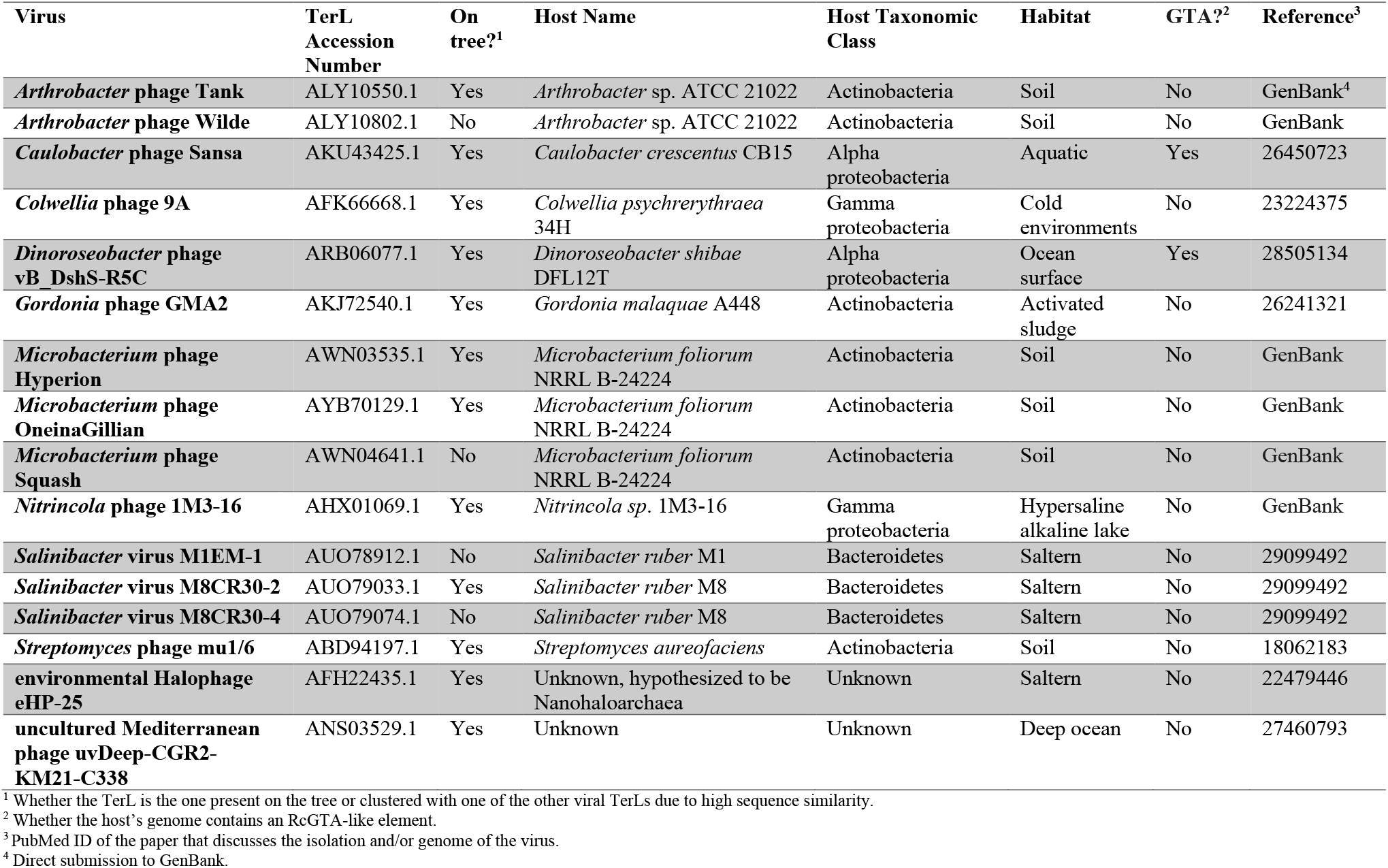
Sixteen viruses with TerLs most closely related to RcGTA-like TerLs.

**Table 2.** Three predicted intact prophages with TerLs closely related to RcGTA-like TerLs.

| Host Name | TerL Accession Number | Predicted prophage coordinates in the host's genome | Host Taxonomic Class | Host Habitat | Reference<sup>1</sup> |
| --- | --- | --- | --- | --- | --- |
| *Zavarzinella formosa* DSM 19928 | WP_020474221.1 | NZ_JH636446.1: 39426-61037 | Planctomycetia | Wetlands | 22740668 |
| *Methylobacterium terrae* 17Sr1-28 | WP_109959484.1 | NZ_CP029553.1: 2863294-2900746 | Alphaproteobacteria | Soil | 31463788 |
| *Thermoactinomyces* sp. DSM 45892 | SDY22851.1 | FNPL01000003.1: 4278-49984 | Bacilli | Not Provided | GenBank<sup>2</sup> |

<sup>1</sup> PubMed ID of the paper that discusses the isolation and/or genome of the host organism
<sup>2</sup> Direct submission to GenBank

Some of the 16 viruses encoding TerLs related to GTA TerLs infect GTA-containing alphaproteobacteria and possess genes that are more closely related to GTA genes than to their homologs in other viruses. For example, *Dinoroseobacter* phage vB_DshS-R5C contains homologs of four putative tail genes of the RcGTA (genes *g12-g15;* Yang et al., 2017). The predicted prophage in *Zavarzinella formosa* DSM 19928 encompasses a homolog of the adaptor gene *(g6)* that is RcGTA-like in amino acid composition. These observations suggest an ongoing exchange and recombination of RcGTA-like genes between viruses and alphaproteobacteria, which likely explains the presence of virus-like TerLs within groups 2 and 3 (**Figure 2**). The TerL phylogeny also indicates that such gene exchange might extend beyond alphaproteobacteria because at least four bacterial TerLs within groups 2 and 3 come from non-alphaproteobacterial genomes (OYV96073, WP_020474221, WP_110156686, and OQX66442). Because the viruses with known hosts infect a wide range of bacteria that live in environments similar to those occupied by GTA-containing alphaproteobacteria (**Table 1**), they either might have an opportunity for gene exchange with viruses that infect GTA-containing bacteria or might be capable of infecting GTA-containing bacteria, in addition to their currently known hosts.

### Two amino acid changes distinguish GTA and viral TerLs

Although viral TerLs that are most closely related to RcGTA-like TerLs have not been experimentally characterized, examination of amino acids that are conserved in RcGTA-like TerLs from large elements (group 3 on **Figure 2**) but not in closely related viral TerLs (group 1 on **Figure 2**), or vice versa, might help pinpoint the changes that are important for the unique packaging strategy of GTAs. We did not identify any amino acids that are conserved in group 1 TerLs but not in group 3 TerLs, but found two amino acids (located at positions 282 and 292 in the RcGTA TerL; RefSeq record WP_031321187) that are conserved in the group 3 TerLs but not in the group 1 TerLs (**Supplementary Table S4** and **Supplementary Figure S1**). In position 292, 99% of the group 3 TerLs, but only 4% of the group 1 TerLs, contain cysteine, whereas 59% of the group 1 TerLs contain threonine. However, given that the threonine to cysteine substitution results in a reduction of the number of carbons per side chain, selection for the reduction in the energetic cost of GTA production (Kogay et al., 2020) cannot be excluded as a driver for this substitution. In position 282, 90% of the group 3 TerLs but no group 1 TerLs contain proline, whereas 64% of the group 1 TerLs but only 6% of the group 3 TerLs contain alanine. Proline contains two more carbons in its side chain than alanine, and therefore, this substitution cannot be selected for energetic cost savings. In TerLs from *bona fide* GTAs of *R. capsulatus* and *Dinoroseobacter shibae*, cysteine is found at position 292 in both proteins, whereas in position 282 the *Dinoroseobacter shibae* TerL has proline and RcGTA TerL has alanine.

Within the TerL protein structure, the two amino acids are located in the nuclease domain (Rao and Feiss, 2015) and lie in close proximity at the opposite ends of a loop that extends towards the translocating DNA (**Supplementary Figure S2** and Figure 3B in Zhao et al. (2013)). The importance of these residues with respect to the functionality of the GTA TerLs remains to be elucidated.

## Discussion

RcGTA is hypothesized to package DNA via a headful mechanism because RcGTA particles encapsidate random pieces of *R. capsulatus’* DNA, which would likely be facilitated by a non-sequence-specific TerL (Casjens et al., 2005; Hynes et al., 2012). Previous experiments support the hypothesis that RcGTA utilizes headful packaging because the packaged DNA fragments have different sequences at the ends (Hynes et al., 2012). The large set of TerL sequences now available allowed us to obtain phylogenetic evidence in support of this hypothesis. Indeed, the RcGTA-like TerLs formed a well-supported, cohesive group within the headful-packaging phage branch in our phylogeny (**Figure 1**).

Previous studies have reported that the RcGTA TerL was most closely related either to the TerLs of T7-like viruses, which use a sequence-specific packaging mechanism (Casjens et al., 2005), or to the TerLs of T4-like viruses, which employ the headful packaging mechanism (Hynes et al., 2012; Lang and Beatty, 2000). However, we found that the RcGTA-like TerLs are more similar to the TerLs of headful-packaging P22-like viruses. This discrepancy is likely due to the vastly expanded set of viral sequences now available in GenBank and the more sensitive search method that we used to identify viral TerL homologs.

In further support of the origin of RcGTA from a virus that employed a headful packaging mechanism, several structural proteins of RcGTA have the highest sequence and secondary structure similarity to the corresponding proteins in viruses that also utilize a headful packaging strategy (Bárdy et al., 2020). Specifically, the tail tape measure protein of RcGTA is homologous to the tail-needle protein of bacteriophage P22, domains of the RcGTA hub and megatron proteins are homologous to their counterparts in bacteriophage T4, and the RcGTA stopper and tail terminator proteins are homologous to those from bacteriophage SPP1 (Bárdy et al., 2020).

We identified TerLs from several viruses and predicted prophages that are phylogenetically closer relatives of the RcGTA TerL than P22-like TerLs. These viruses and predicted prophages either infect alphaproteobacteria or at least are found in the same environments as GTA-containing alphaproteobacteria. Because the specific mechanisms of headful packaging differ among phages (Bhattacharyya and Rao, 1993; Casjens et al., 1992, 2004), experimental characterization of packaging in viruses that are closely related to GTAs could offer further insight into the origin of the GTA TerLs and their packaging mechanism.

The TerL phylogeny presented here supports the single origin of the RcGTA head-tail cluster in alphaproteobacteria because large RcGTA-like elements grouped together. With the newfound support for a single origin of RcGTA-like TerLs from a TerL of a headful-packaging phage, we propose that a headful-packaging TerL in the RcGTA ancestor underwent key changes that resulted in the switch from packaging the GTA genome to packaging random, small pieces of the host genome (Hynes et al., 2012) with a substantially lower density of DNA in the capsid (Bárdy et al., 2020). The selection for reduced energy cost of GTA protein production that apparently occurred after the origin of RcGTA-like elements in alphaproteobacteria (Kogay et al., 2020) makes it challenging to pinpoint amino acid changes in GTA TerLs that contribute to this transition.

The loss of specificity for the GTA genome and the reduction in the DNA packaging density rule out self-propagation of the GTA genome mediated by virions. Strikingly, the RcGTA genes appear to be actively precluded from packaging, being the least frequently packaged region in the alphaproteobacterial genome that is incorporated into the GTA particles (Hynes et al., 2012). The mechanism behind this exclusion is unknown, one possibility being that intensive expression of the RcGTA genes interferes with their packaging (Hynes et al., 2012).

In addition to TerL, other GTA proteins that are involved in DNA packaging, including TerS and portal, might contribute to the unique DNA packaging features of RcGTA. In particular, the RcGTA gene *g1*, which is adjacent to the *terL* gene *(g2)* in the RcGTA genome, has been recently shown to encode TerS (Sherlock et al., 2019). Sherlock et al. (2019) demonstrated that the RcGTA TerS binds non-specifically to DNA with low affinity due to the absence of a specific DNA-binding domain and the retention of non-specific DNA binding activity. A TerS protein with altered DNA-binding characteristics and a modified headful TerL might together underlie the random packaging that is characteristic of RcGTA.

## Author Contributions

E.E., O.Z., Y.I.W. and E.V.K. designed the study. E.E. collected data. E.E. and R.K. performed the analyses. E.E., O.Z., Y.I.W., R.K. and E.V.K. interpreted the results. E.E. and O.Z. wrote the initial draft of the manuscript. E.E., O.Z., Y.I.W., R.K. and E.V.K. revised the manuscript.

## Acknowledgements

We thank Zhengshuang Hua for insightful discussions and help with using Dartmouth computing facilities. This work was supported by the following awards from Dartmouth College to E.E.: Sophomore Research Scholarship, James O. Freedman Presidential Scholarship, Thomas B. Roos Memorial Fund Fellowship and a Kaminsky Undergraduate Research Award. Additionally, this work was supported by an Intramural Research and Training Award from the National Institutes of Health to E.E., by the Simons Foundation Investigator in Mathematical Modeling of Living Systems award #327936 to O.Z., by the National Science Foundation award DEB-1551674 to O.Z., and by the Intramural Research Program of the U.S. National Institutes of Health (National Library of Medicine) to Y.I.W. and E.V.K.

## Data Availability

The following data is provided in our **Supplementary Dataset** available via FigShare (https://doi.org/10.6084/m9.figshare.12191691): GenBank accession numbers of the 254,382 RcGTA TerL protein homologs that are taxonomically assigned to bacteria, archaea, or viruses and likely include both ATPase and nuclease domains, GenBank accession numbers of the 252,614 RcGTA TerL protein homologs that are represented by 11,051 TerLs in the tree in **Figure 1**, GenBank accession numbers of the 11,051 amino acid sequences used for reconstruction of the tree in **Figure 1**, alignment of 11,051 TerL amino acid sequences (untrimmed and trimmed), phylogenetic tree of 11,051 TerLs shown in **Figure 1**, alignment of amino acid sequences of 616 TerLs from the subtree shown in **Figure 2**, alignments and phylogenetic trees of 342 and 346 TerLs used in IQ-TREE phylogenetic reconstructions. All alignments are in FASTA format and all phylogenetic trees are in NEWICK format.

## Supplementary Tables

(in a separate file in Excel format)

**Supplementary Table S1:** List of the 20 profiles used as queries for PSI-BLAST searches to find TerL homologs.

**Supplementary Table S2:** List of 73 viral TerLs of experimentally determined packaging mechanism that were identified in our dataset of 252,614 TerL homologs.

**Supplementary Table S3:** GTA-Hunter classifications of 604 bacterial and archaeal TerLs from the RcGTA-containing subtree of TerL phylogeny and PHASTER predictions of prophage regions surrounding the virus-like TerLs.

**Supplementary Table S4:** The amino acids at the sites homologous to the positions 282 and 292 in the RcGTA TerL (WP_031321187) in 616 bacterial and viral TerLs from the subtree shown in Figure 2.

## Supplementary Figures

**Supplementary Figure S1.**
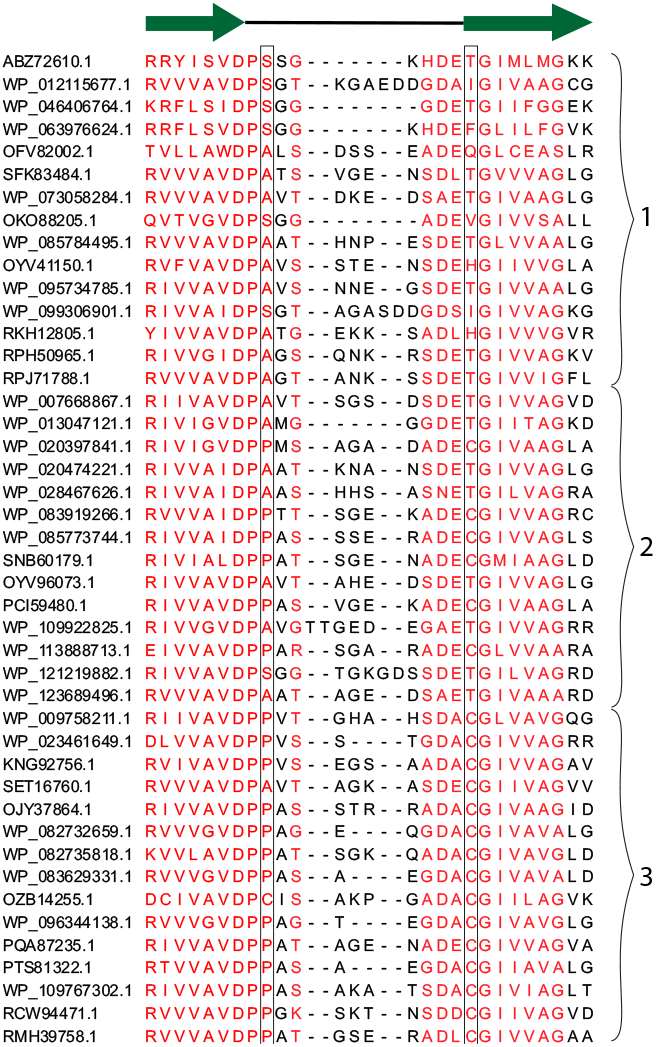
Multiple sequence alignment of the TerL region with two conserved sites that differentiate GTA and viral TerLs. The shown selection of representative sequences is designated by their RefSeq or GenBank identifiers. The selected sequences representing the three groups from **Figure 2** (marked by curly braces). The secondary structure of the region is shown above the alignment, where green arrows designate beta strands and the black line indicates a random coil. Alignment sites colored in red contain at least 2 bits of information. The two conserved differentiating sites are outlined by rectangles. The amino acids found in these sites in the full dataset are provided in **Supplementary Table S4**. The depicted region of the protein is highlighted on the TerL protein structure shown in **Supplementary Figure S2**).

**Supplementary Figure S2.**
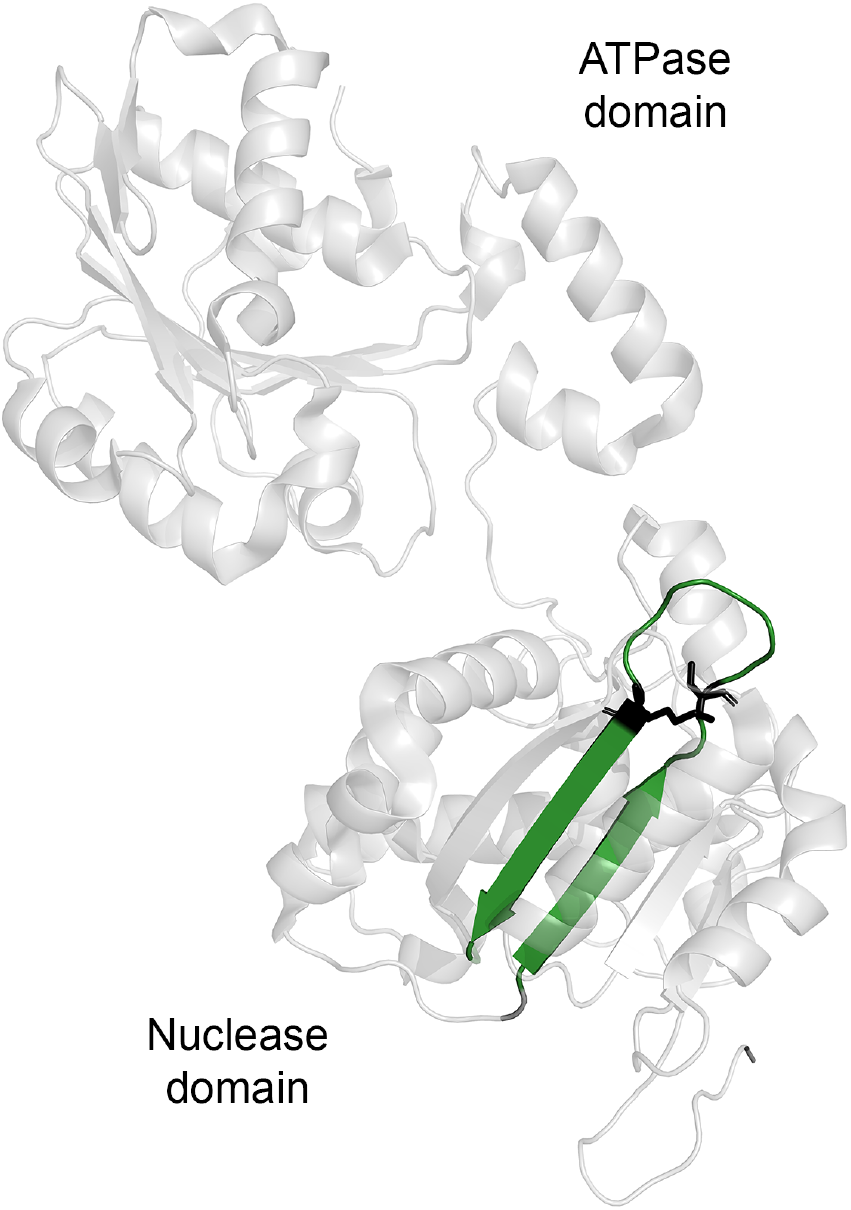
Locations of the two conserved differentiating sites on the protein structure of the TerL from *Shigella* phage Sf6 (PDB ID 4IDH) (Zhao et al., 2013). The side chains of the two amino acids corresponding to the differentiating sites (S266 and K273) are shown in black. The two beta strands and a random coil region shown in **Supplementary Figure S1** are colored in green. The rest of the structure is shown at reduced opacity for presentation purposes.

## References

Altschul, S.F., Madden, T.L., Schäffer, A.A., Zhang, J., Zhang, Z., Miller, W., and Lipman, D.J. (1997). Gapped BLAST and PSI-BLAST: a new generation of protein database search programs. Nucleic Acids Res 25, 3389–3402.

Arndt, D., Grant, J.R., Marcu, A., Sajed, T., Pon, A., Liang, Y., and Wishart, D.S. (2016). PHASTER: a better, faster version of the PHAST phage search tool. Nucleic Acids Res 44, W16–21.

Bárdy, P., Füzik, T., Hrebík, D., Pantůček, R., Thomas Beatty, J., and Plevka, P. (2020). Structure and mechanism of DNA delivery of a gene transfer agent. Nat Commun 11, 3034.

Bhattacharyya, S.P., and Rao, V.B. (1993). A novel terminase activity associated with the DNA packaging protein gp17 of bacteriophage T4. Virology 196, 34–44.

Casjens, S.R. (2011). The DNA-packaging nanomotor of tailed bacteriophages. Nat Rev Microbiol 9, 647–657.

Casjens, S.R., and Gilcrease, E.B. (2009). Determining DNA Packaging Strategy by Analysis of the Termini of the Chromosomes in Tailed-Bacteriophage Virions. Methods Mol Biol 502, 91–111.

Casjens, S., Sampson, L., Randall, S., Eppler, K., Wu, H., Petri, J.B., and Schmieger, H. (1992). Molecular genetic analysis of bacteriophage P22 gene 3 product, a protein involved in the initiation of headful DNA packaging. J Mol Biol 227, 1086–1099.

Casjens, S., Winn-Stapley, D.A., Gilcrease, E.B., Morona, R., Kühlewein, C., Chua, J.E.H., Manning, P.A., Inwood, W., and Clark, A.J. (2004). The chromosome of *Shigella flexneri* bacteriophage Sf6: complete nucleotide sequence, genetic mosaicism, and DNA packaging. J Mol Biol 339, 379–394.

Casjens, S.R., Gilcrease, E.B., Winn-Stapley, D.A., Schicklmaier, P., Schmieger, H., Pedulla, M.L., Ford, M.E., Houtz, J.M., Hatfull, G.F., and Hendrix, R.W. (2005). The Generalized Transducing *Salmonella* Bacteriophage ES18: Complete Genome Sequence and DNA Packaging Strategy. J Bacteriol 187, 1091–1104.

Ding, H., Grüll, M.P., Mulligan, M.E., Lang, A.S., and Beatty, J.T. (2019). Induction of *Rhodobacter capsulatus* Gene Transfer Agent Gene Expression Is a Bistable Stochastic Process Repressed by an Extracellular Calcium-Binding RTX Protein Homologue. J Bacteriol 201.

Edgar, R.C. (2004). MUSCLE: a multiple sequence alignment method with reduced time and space complexity. BMC Bioinformatics 5, 113.

Edgar, R.C. (2010). Search and clustering orders of magnitude faster than BLAST. Bioinformatics 26, 2460–2461.

Ester, M., Kriegel, H.-P., Sander, J., and Xu, X. (1996). A density-based algorithm for discovering clusters in large spatial databases with noise. In Proceedings of the Second International Conference on Knowledge Discovery and Data Mining, (Portland, Oregon: AAAI Press), pp. 226–231.

Fogg, P.C.M. (2019). Identification and characterization of a direct activator of a gene transfer agent. Nat Commun 10, 595.

Fogg, P.C.M., Westbye, A.B., and Beatty, J.T. (2012). One for all or all for one: heterogeneous expression and host cell lysis are key to gene transfer agent activity in *Rhodobacter capsulatus*. PLOS ONE 7, e43772.

Fokine, A., and Rossmann, M.G. (2014). Molecular architecture of tailed double-stranded DNA phages. Bacteriophage 4, e28281.

Fu, Y., Macleod, D.M., Rivkin, R.B., Chen, F., Buchan, A., and Lang, A.S. (2010). High diversity of *Rhodobacterales* in the subarctic North Atlantic Ocean and gene transfer agent protein expression in isolated strains. Aquat Microb Ecol 59, 283–293.

Grull, M., Mulligan, M., and Lang, A. (2018). Small extracellular particles with big potential for horizontal gene transfer: membrane vesicles and gene transfer agents. FEMS Microbiol Lett 365, fny192.

Henikoff, S., and Henikoff, J.G. (1993). Performance evaluation of amino acid substitution matrices. Proteins 17, 49–61.

Henikoff, S., and Henikoff, J.G. (1994). Position-based sequence weights. J Mol Biol 243, 574–578.

Hoang, D.T., Chernomor, O., von Haeseler, A., Minh, B.Q., and Vinh, L.S. (2018). UFBoot2: Improving the Ultrafast Bootstrap Approximation. Mol Biol Evol 35, 518–522.

Hynes, A.P., Mercer, R.G., Watton, D.E., Buckley, C.B., and Lang, A.S. (2012). DNA packaging bias and differential expression of gene transfer agent genes within a population during production and release of the *Rhodobacter capsulatus* gene transfer agent, RcGTA. Mol Microbiol 85, 314–325.

Hynes, A.P., Shakya, M., Mercer, R.G., Grüll, M.P., Bown, L., Davidson, F., Steffen, E., Matchem, H., Peach, M.E., Berger, T., et al. (2016). Functional and Evolutionary Characterization of a Gene Transfer Agent’s Multilocus “Genome.” Mol Biol Evol 33, 2530–2543.

Kalyaanamoorthy, S., Minh, B.Q., Wong, T.K.F., von Haeseler, A., and Jermiin, L.S. (2017). ModelFinder: fast model selection for accurate phylogenetic estimates. Nat Methods 14, 587–589.

Katoh, K., and Standley, D.M. (2013). MAFFT Multiple Sequence Alignment Software Version 7: Improvements in Performance and Usability. Mol Biol Evol 30, 772–780.

Kogay, R., Neely, T.B., Birnbaum, D.P., Hankel, C.R., Shakya, M., and Zhaxybayeva, O. (2019). Machine-Learning Classification Suggests That Many Alphaproteobacterial Prophages May Instead Be Gene Transfer Agents. Genome Biol Evol 11, 2941–2953.

Kogay, R., Wolf, Y.I., Koonin, E.V., and Zhaxybayeva, O. (2020). Selection for Reducing Energy Cost of Protein Production Drives the GC Content and Amino Acid Composition Bias in Gene Transfer Agents. MBio 11, e01206–20.

Lang, A.S., and Beatty, J.T. (2000). Genetic analysis of a bacterial genetic exchange element: The gene transfer agent of *Rhodobacter capsulatus*. Proc Natl Acad Sci USA 97, 859–864.

Lang, A.S., Westbye, A.B., and Beatty, J.T. (2017). The Distribution, Evolution, and Roles of Gene Transfer Agents in Prokaryotic Genetic Exchange. Annu Rev Virol 4, 87–104.

Lu, S., Wang, J., Chitsaz, F., Derbyshire, M.K., Geer, R.C., Gonzales, N.R., Gwadz, M., Hurwitz, D.I., Marchler, G.H., Song, J.S., et al. (2020). CDD/SPARCLE: the conserved domain database in 2020. Nucleic Acids Res 48, D265–D268.

McDaniel, L.D., Young, E., Delaney, J., Ruhnau, F., Ritchie, K.B., and Paul, J.H. (2010). High Frequency of Horizontal Gene Transfer in the Oceans. Science 330, 50.

Merrill, B.D., Ward, A.T., Grose, J.H., and Hope, S. (2016). Software-based analysis of bacteriophage genomes, physical ends, and packaging strategies. BMC Genomics 17, 679.

Nagao, N., Yamamoto, J., Komatsu, H., Suzuki, H., Hirose, Y., Umekage, S., Ohyama, T., and Kikuchi, Y. (2015). The gene transfer agent-like particle of the marine phototrophic bacterium *Rhodovulum sulfidophilum*. Biochem Biophys Rep 4, 369–374.

Nguyen, L.-T., Schmidt, H.A., von Haeseler, A., and Minh, B.Q. (2015). IQ-TREE: A Fast and Effective Stochastic Algorithm for Estimating Maximum-Likelihood Phylogenies. Mol Biol Evol 32, 268–274.

Philosof, A., Yutin, N., Flores-Uribe, J., Sharon, I., Koonin, E.V., and Béjà, O. (2017). Novel Abundant Oceanic Viruses of Uncultured Marine Group II Euryarchaeota. Curr Biol 27, 1362–1368.

Price, M.N., Dehal, P.S., and Arkin, A.P. (2010). FastTree 2 – Approximately Maximum-Likelihood Trees for Large Alignments. PLOS ONE 5, e9490.

Rao, V.B., and Feiss, M. (2008). The Bacteriophage DNA Packaging Motor. Annu Rev Genet 42, 647–681.

Rao, V.B., and Feiss, M. (2015). Mechanisms of DNA Packaging by Large Double-Stranded DNA Viruses. Annu Rev Virol 2, 351–378.

Schrödinger, LLC (2020). The PyMOL Molecular Graphics System, Version 2.4.

Shakya, M., Soucy, S.M., and Zhaxybayeva, O. (2017). Insights into origin and evolution of α-proteobacterial gene transfer agents. Virus Evol 3, vex036.

Sherlock, D., Leong, J.X., and Fogg, P.C.M. (2019). Identification of the First Gene Transfer Agent (GTA) Small Terminase in *Rhodobacter capsulatus* and Its Role in GTA Production and Packaging of DNA. J Virol 93, e01328–19.

Söding, J. (2005). Protein homology detection by HMM–HMM comparison. Bioinformatics 21, 951–960.

Sokal, R.R., and Michener, C.D. (1958). A statistical method for evaluating systematic relationships. Univ Kansas Sci Bull 38, 1409–1438.

Solioz, M., Yen, H.C., and Marris, B. (1975). Release and uptake of gene transfer agent by *Rhodopseudomonas capsulata*. J Bacteriol 123, 651–657.

Steinegger, M., and Söding, J. (2017). MMseqs2 enables sensitive protein sequence searching for the analysis of massive data sets. Nat Biotechnol 35, 1026–1028.

Westbye, A.B., O’Neill, Z., Schellenberg-Beaver, T., and Beatty, J.T. (2017a). The *Rhodobacter capsulatus* gene transfer agent is induced by nutrient depletion and the RNAP omega subunit. Microbiology 163, 1355–1363.

Westbye, A.B., Beatty, J.T., and Lang, A.S. (2017b). Guaranteeing a captive audience: coordinated regulation of gene transfer agent (GTA) production and recipient capability by cellular regulators. Curr Opin Microbiol 38, 122–129.

Whelan, S., and Goldman, N. (2001). A general empirical model of protein evolution derived from multiple protein families using a maximum-likelihood approach. Mol Biol Evol 18, 691–699.

Wolf, Y.I., Kazlauskas, D., Iranzo, J., Lucía-Sanz, A., Kuhn, J.H., Krupovic, M., Dolja, V.V., and Koonin, E.V. (2018). Origins and Evolution of the Global RNA Virome. MBio 9, e02329–18.

Yang, Z. (1994). Maximum likelihood phylogenetic estimation from DNA sequences with variable rates over sites: Approximate methods. J Mol Evol 39, 306–314.

Yang, Y., Cai, L., Ma, R., Xu, Y., Tong, Y., Huang, Y., Jiao, N., and Zhang, R. (2017). A Novel Roseosiphophage Isolated from the Oligotrophic South China Sea. Viruses 9, 109.

Yutin, N., Makarova, K.S., Gussow, A.B., Krupovic, M., Segall, A., Edwards, R.A., and Koonin, E.V. (2018). Discovery of an expansive bacteriophage family that includes the most abundant viruses from the human gut. Nat Microbiol 3, 38–46.

Zhao, H., Christensen, T.E., Kamau, Y.N., and Tang, L. (2013). Structures of the phage Sf6 large terminase provide new insights into DNA translocation and cleavage. Proc Natl Acad Sci USA 110, 8075–8080.

Zimmermann, L., Stephens, A., Nam, S.-Z., Rau, D., Kübler, J., Lozajic, M., Gabler, F., Söding, J., Lupas, A.N., and Alva, V. (2018). A Completely Reimplemented MPI Bioinformatics Toolkit with a New HHpred Server at its Core. J Mol Biol 430, 2237–2243.

